# Real-time heart rate in the wild: remote collection of cardiac data in baboons using a low-power Bluetooth and LoRaWAN system

**DOI:** 10.64898/2026.04.17.719194

**Authors:** Erin S. Person, Catherine R. Andreadis, Annabelle G. Beaton, Ann N. Namunyak, Edward Kariuki, Paul Solheim, Amanda Taylor, Peter Leimgruber, Rosana N. Moraes, Paul A. Iaizzo, Jenny Tung, Herman Pontzer, Mercy Y. Akinyi, Susan C. Alberts, Timotheus J. van Dam, Timothy G. Laske, Elizabeth A. Archie

**Author notes:** Corresponding author: Erin Person.

## Abstract

1. Cardiac rate and rhythm reveal how animals adapt physiologically to day-to-day challenges, with consequences for health and fitness. However, these data remain difficult to collect in wild animals, despite their relevance for individual health and fitness.
2. Here, we present a system for collecting and transmitting long-term, fine-scaled physiological data in wild animals. We implanted Bluetooth-enabled cardiac and physiological monitor devices in three wild adult female baboons in the Amboseli ecosystem in Kenya and paired these devices with collars that enabled remote data downloads over long-range wide area network (LoRaWAN).
3. The system performed well over >10 months, providing the first long-term cardiac data in wild primates. The baboons showed strong circadian patterns in heart rate, heart rate variability, and activity. We also present data on one female who left her social group for unknown reasons; while alone she exhibited higher heart rate variability, lower activity, and evidence of disrupted sleep.
4. In sum, physiologgers paired with low-energy methods of remote data retrieval are powerful tools for investigating physiology in wild animals on timescales that extend over many months, with minimal disruption to their behavior.

## Introduction

Fine-grained, temporally resolved data on cardiac rate and rhythm are valuable for understanding animal health and responses to environmental change (Gaidica & Dantzer, 2020; Hawkes et al., 2021). High-resolution data on heart rate (HR) and heart rate variability (HRV) have revealed natural patterns of energy expenditure and emotional arousal during stressors in multiple species (e.g., Brandl et al., 2025; Wascher, 2021). Such data have already been used to manage cardiovascular disease in zoo-housed great apes (Kutinsky et al., 2023).

Despite their potential, collecting data on real-time changes in cardiac metrics in wild animals is challenging. Among the best available tools are subcutaneous physiologgers, called insertable cardiac monitors (ICMs), which record electrical signals from the heart (Kutinsky et al., 2023; Laske et al., 2011, 2017, 2018; Leimgruber et al., 2023; Moraes et al., 2021). While ICMs can collect precise, high-resolution data on cardiac metrics, most ICMs require researchers to recapture the animals to recover the data (Forin-Wiart et al., 2019). Recapture is difficult or impossible for many species, leading to a high risk of data loss. While some ICMs use telemetry or cellular/satellite networks to retrieve data, these options are power-hungry, which constrains ICM battery life (Brandl et al., 2025; Wild, van Schalkwyk, et al., 2023; Williams et al., 2021).

One solution lies in low power data transmission strategies such as Bluetooth Low-Energy (BLE) and long-range wide area networks (LoRaWAN). BLE transmission distances are often short but their power draw is minimal, which is ideal for devices with small batteries like ICMs. When paired with a LoRaWAN-enabled collar, which can hold larger batteries, data can be transmitted over long distances (Devalal & Karthikeyan, 2018). LoRaWAN systems are already in use in livestock care (Perea et al., 2025) and have more recently been applied to wildlife monitoring (LoRa Alliance™ Strategy Committee, 2018). However, to our knowledge, it has not been implemented for physiologger data in wild animals (Antoine-Santoni et al., 2018; Gauld et al., 2023).

We present a novel system to record and transmit real-time data on HR, HRV, body temperature, and physical activity in wild baboons, using ICMs with BLE capability paired with LoRaWAN-enabled collars. We piloted this system in a well-studied population of wild baboons living in a remote area in Kenya. The system satisfied four requirements: (1) continuous data collection over several months to capture cardiac responses to common and rare events; (2) data downloads at distances >20 m to minimize disruption to the animals; (3) data uploads to a cloud server from a remote site with unreliable cellular service; and (4) reliable performance in the absence of permanent network infrastructure at the site (e.g., tower with a LoRaWAN gateway; Gauld et al., 2023).

Here we describe the system and present information on the speed, success, and range of data transmission. We also report preliminary data on HR and HRV in wild baboons and share a case study of an older, pregnant, female baboon who left her social group and spent nine days alone—an event that may have been linked to illness and was likely very stressful. We show that BLE and LoRaWAN can work in tandem to transmit physiologger data over tens to hundreds of meters, enabling physiological data collection in the wild.

## Methods

### Study population and subjects

Our subjects were three adult female baboons studied by the Amboseli Baboon Research Project (ABRP) in southern Kenya (Alberts & Altmann, 2012). The population is composed of admixed baboons with dominant *Papio cynocephalus* ancestry combined with *P*.*anubis* (Vilgalys et al., 2022). Our subjects lived in a single social group of individually recognized, habituated baboons. All three had been observed from birth and were aged 4.9, 12.0, and 22.2 years at ICM implantation. Experienced observers visited the group every 2-4 days to collect group censuses, record behavioral data, download ICM data, and monitor health.

### Hardware

Figure 1 shows an overview of the system. We used LINQ II™ ICMs developed for human clinical use and donated by Medtronic Inc. (Fig. 2A; Medtronic Inc., Minneapolis, MN, USA). The LINQ II™ is lightweight and has an expected battery life of 4.5 years (mass=3.4 g; dimensions=45.1 x 8.0 x 4.2 mm). The ICMs were programmed to record and transmit 50 bytes of data at 5-minute intervals (data logs). These data included: (1) median HR in beats per minute (bpm); (2) HRV as the root mean square of successive differences between beats (RMSSD) in milliseconds; (3) activity level from an on-board triaxial accelerometer, encoded on a unitless scale from 0 to 255; (4) body temperature in °C once per hour.

**Figure 1.**
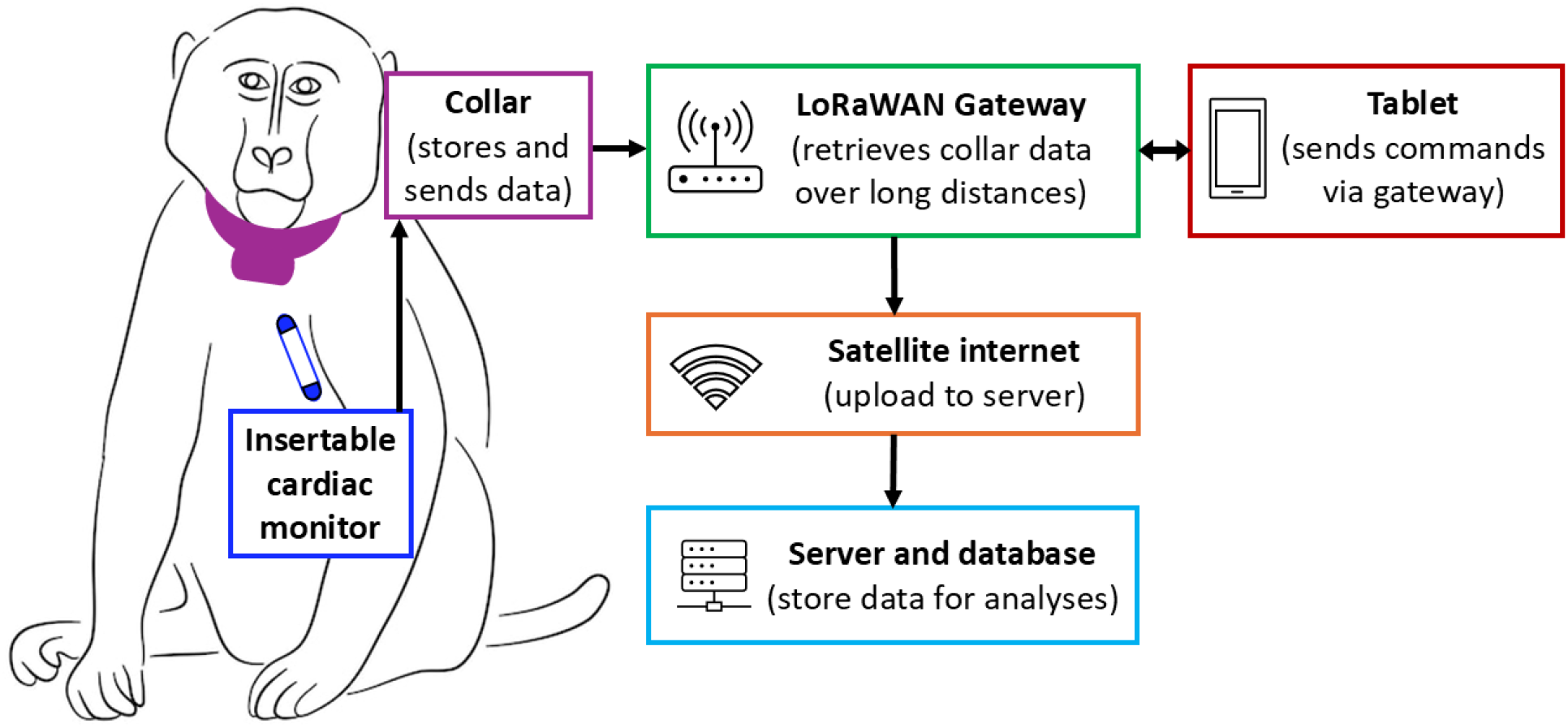
ICM and remote data download system. A LINQ II™ ICM was deployed subcutaneously (dark blue) over the left pectoral muscle of a female baboon (Andreadis et al., 2026). ICM data were transmitted every 5 minutes via BLE to a lightweight, battery-powered collar (purple) that stored the data. Observers downloaded collar data using a “mobile download unit”, which included a mobile LoRaWAN gateway (green) with an omni-directional antenna and an Android tablet (red) connected to satellite internet (orange). Using the tablet, an observer sent requests to the gateway to download logs, which were received and transmitted to a cloud server (light blue). Arrows indicate the flow of data through the system. The collar advertised a status update and received messages from the gateway every 15 minutes to save battery.

**Figure 2.**
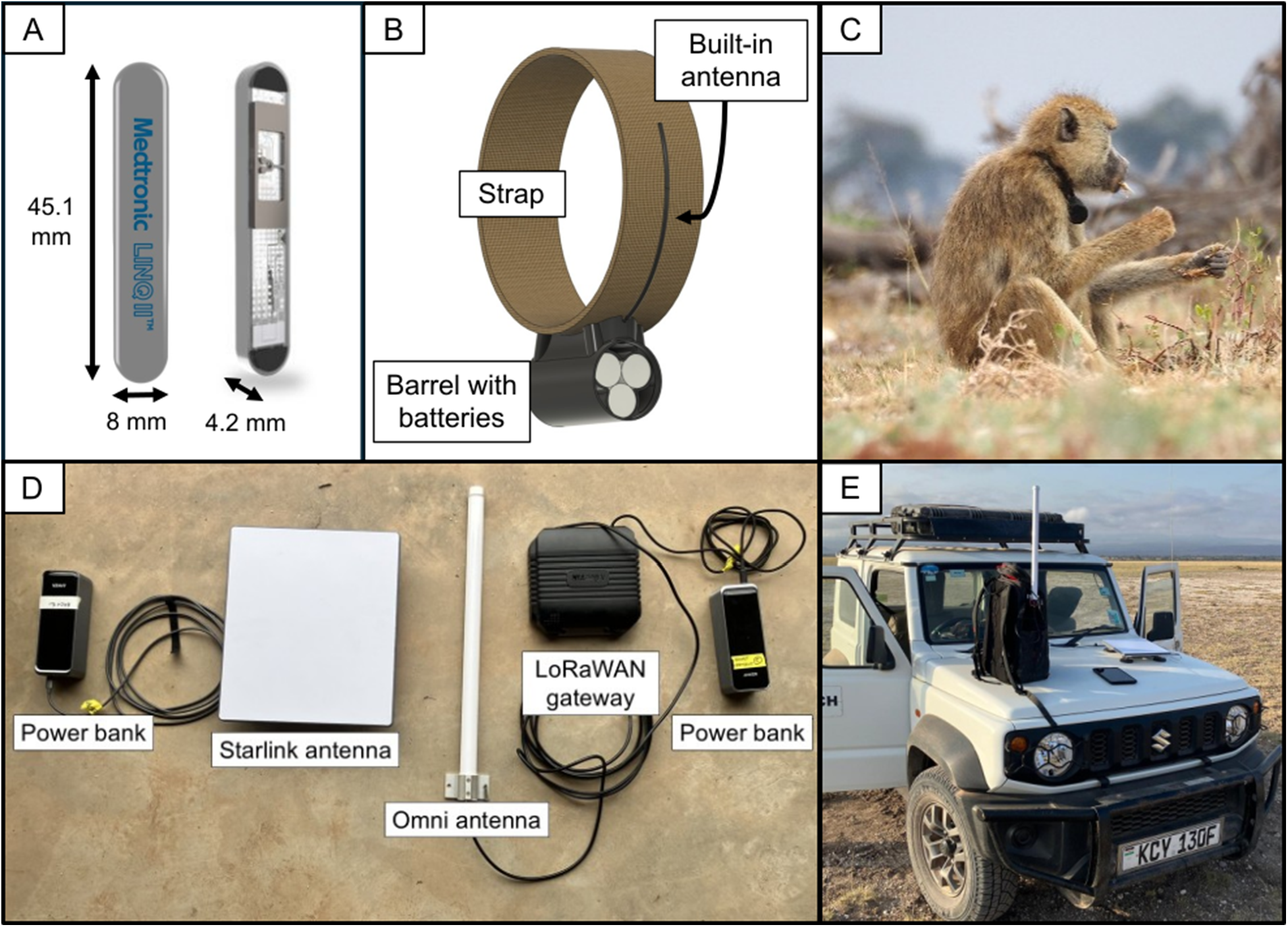
Hardware. (A) Medtronic LINQ II™ ICM. (B) Smart Parks collar with 3D-printed case containing the electronic components and built-in LoRaWAN wire antenna. (C) Baboon wearing a collar (photo credit: Susan Alberts) (D) Components of the mobile download unit (excluding the tablet). (E) The mobile download unit deployed on a vehicle (Starlink antenna magnetically mounted to the vehicle’s hood; gateway and omni antennas in the backpack).

The LoRaWAN-enabled collars were developed by Smart Parks (Figs. 2B,C; Utrecht, The Netherlands) and consisted of a 3D-printed acrylonitrile styrene acrylate case (35 x 64 mm) and biothane strap (22 mm x 406 mm; mass=120 g, <1.25% of the females’ body mass). The electronics included a printed circuit board (16 MB flash storage), external LoRaWAN wire antenna, and internal BLE antenna, powered by 3 AA batteries (LS14500 cell, Saft, Levallois-Perret, France). The LoRaWAN gateway was a LtAP LR8G LTE6 gateway kit (Mikrotik, Riga, Latvia; Fig. 2D) with a 6.5 dBi omni-directional antenna (868 antenna kit, Mikrotik). We used a Galaxy Active3 Tab tablet (Samsung, Seoul, South Korea) connected to satellite internet via a Starlink Mini Terminal (Starlink, Hawthorne, CA, USA). The gateway and Starlink antenna were powered by Anker Prime Power Banks (250W, Changsha, Hunan, China). We used a magnetic mount to attach the Starlink antenna to the vehicle (NovaKits, third party retailer on Amazon.com, Fig. 2E). We carried the gateway, omni antenna, and power banks in a backpack.

### Software

To pair the ICMs to the collars and set the collar advertisement frequency, we used the Smart Parks Connect app (v2.10.2) for Android created by the collar developer (TvD). We used ChirpStack, an open source LoRaWAN network server (v4, Brocaar 2025) to send and receive messages from the gateway, send commands to the collars, and monitor packets of data logs received from the collars. We used Node-RED (v3.1.3), an open-source software, to collect, transform, and visualize the packets, which were then sent to an InfluxDB Database (v3, InfluxData Inc), an open-source time series database. Grafana (v10.3.1, Grafana Labs), a web app for data analysis and visualization, was used to query the InfluxDB database, display the data in graphical form, and export the data for further analysis.

### ICM insertion

ICMs were inserted subcutaneously following Andreadis et al. (2026; see Supplementary Methods).

### Permits

All research was approved by the IACUC at Duke University (A003-24-01), University of Notre Dame (25-01-9041), and the Ethics Council of the Max Planck Society (2022_29) and adhered to all the laws and guidelines of Kenya, including permits from the Wildlife Research Training Institute (WRTI/RPC/2025-226599), the National Commission for Science, Technology, and Innovation (NACOSTI/P/25/4181604), and the National Environmental Management Authority (NEMA/AGR/198/2024).

### Tests and analyses

To measure data download speeds and data packet loss, we examined 12 weeks of data for each baboon. To test the effective range of our mobile gateway system, we moved the gateway progressively farther from the collared females and used GPS locations to measure the distance between them. We noted the farthest distance at which we could still detect a signal from the collars. We further tested whether downloading data from two or more collars simultaneously affected the success of the download by monitoring the number of lost data packets (i.e., data sent from the collar but not received by the gateway).

## Results

Table 1 shows the identity, implantation date, age, and body mass for each baboon. All baboons tolerated the ICMs well and showed no signs of infection following surgery. At the time of writing (February 2026), two ICM/collars sets remain functional; the third belonged to the female HUC who disappeared from her social group six months after implantation (see Case Study in Supporting Information).

**Table 1.** Study subject, implantation date, age, and body mass at implantation.

| Subject | Date | Age (years) | Mass (kg) |
| --- | --- | --- | --- |
| HUC | 06-Mar-25 | 22.2 | 14.0 |
| GAG | 03-Mar-25 | 4.9 | 9.6 |
| FAU | 03-Mar-25 | 12.0 | 13.5 |

### Performance of the mobile download system

Table 2 shows performance metrics of the remote download system. 24 hours of data (~98.4 KB) could be downloaded in about 2.8 minutes. We typically downloaded 4-8 days of data from an animal at a time, which required 11-25 minutes per individual (2.8 minutes x 4-8 days=11-25 minutes). As expected, some data packets were lost between the collar and the LoRaWAN gateway. When we noted missing packets, we re-initiated the download (our packet loss calculations include these re-initiated downloads; Table 2). Packet loss varied with the distance from the gateway antenna to the target animal and with the presence of obstructions (e.g., vegetation, hills). Downloading from multiple collars simultaneously resulted in large increases of packet loss due to constraints of the LoRaWAN gateway. Downloads were most consistent when the gateway antenna was <100 m from the baboon. The farthest distance we were able to receive a collar’s status update was 975 m.

**Table 2.** Performance metrics of the remote download system.

| Download rate | Collar storage | Mean daily packet loss |
| --- | --- | --- |
| 35 KB per minute | 16 MB<br>(~166 days of data) | 5.2%<br>(15±21 of 288 data logs lost; range=1-175) |

### Physiological data recorded by the ICMs

Table 3 summarizes 5-minute HR, HRV, and activity scores, and 1-hour body temperature for all subjects combined and for each subject separately. Figure 3 visualizes these values for the female “GAG” between 21-28 August, 2025. HR displayed a circadian rhythm with a diurnal peak in the afternoon (~14:00-16:00) and a trough at ~05:00. HRV also followed a circadian pattern, peaking in the early morning when the baboon became active. Activity was highest during daylight (06:30-18:30), when the baboons were awake and moving, and lowest at night (Fig. 3, Fig. S1). Typical daytime activities (e.g., walking, foraging, socializing) registered as high activity scores on the unitless activity scale; median daytime activity was 212 while median nighttime activity was 27 (Fig. 3, Fig. S1). Body temperature did not display a strong circadian pattern.

**Table 3.** Physiological metrics across and within all subjects between March-August 2025.

|  |  | All subjects | HUC | GAG | FAU |
| --- | --- | --- | --- | --- | --- |
| Heart Rate (bpm) | Mean ± SD | 99.94±22.08 | 91.93±21.26 | 105.3±23.55 | 101.9±18.89 |
|  | Range | 41.1-176.5 | 41.1-171.4 | 55.56-176.5 | 50-176.5 |
| Heart rate variability (ms) | Mean ± SD | 4.639±3.484 | 4.527±3.138 | 6.179±3.789 | 3.147±2.698 |
|  | Range | 0-48.3 | 0-48.3 | 0-28.3 | 0-17 |
| Activity score | Mean ± SD | 115.6±92.01 | 109.9±90.6 | 114.8±92.54 | 121.7±92.39 |
|  | Range | 14-255 | 14-255 | 15-255 | 15-255 |
| Body temperature (°C) | Mean ± SD | 36.88±0.8705 | 36.95±0.977 | 36.5±0.7105 | 37.21±0.7575 |
|  | Range | 31.8-40 | 31.8-39.5 | 33.5-38.8 | 33.5-40 |

**Figure 3.**
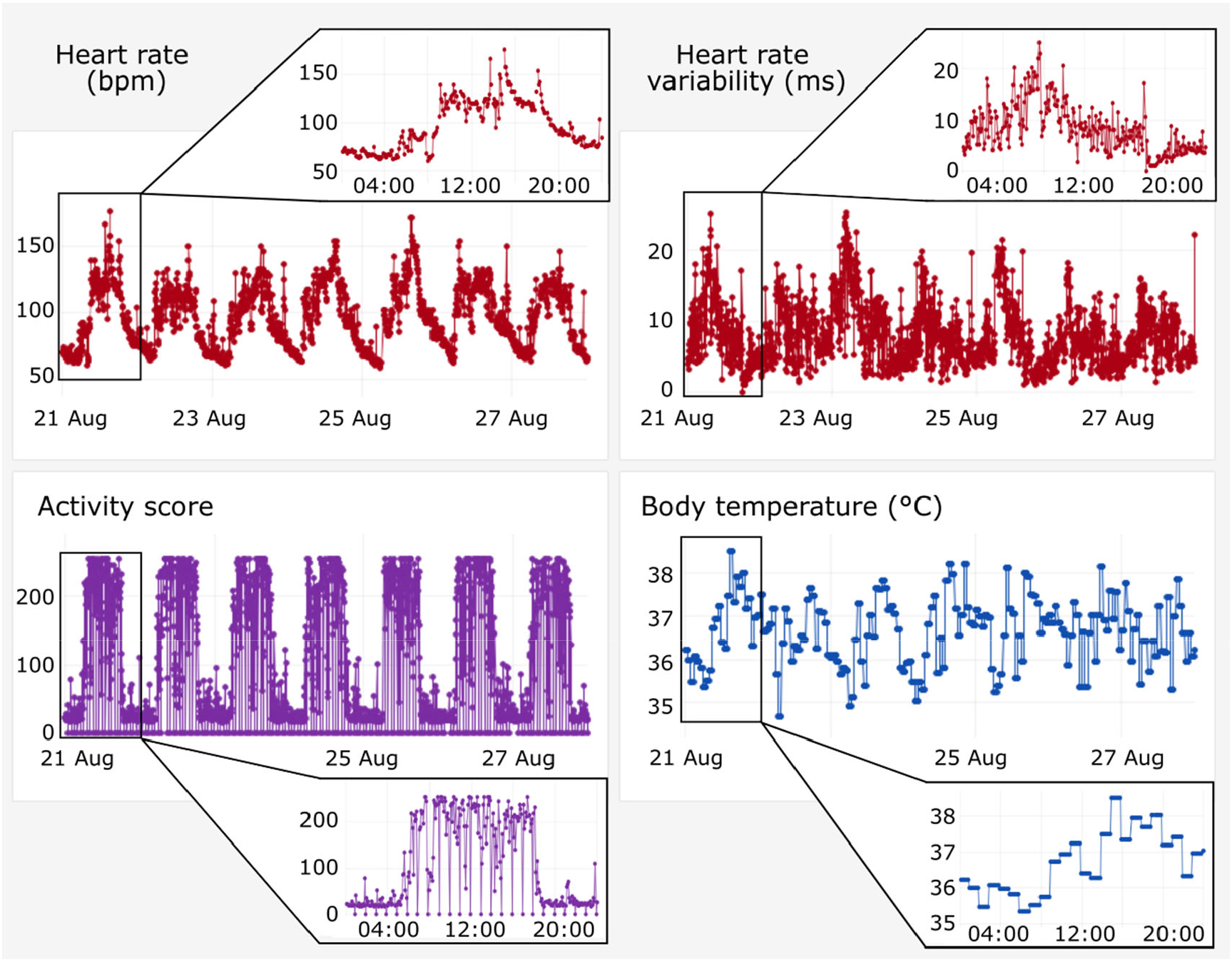
5-minute HR, HRV, and activity, and 1-hour body temperature for the female “GAG” over a representative 7-day period. X-axes show date and time; y-axes show HR, HRV, activity score, and body temperature. Data were collected between 21-28 August; insets show data for a 24-hour period on August 21. The hourly 0 values for activity score were due to a programming error in the ICM; these values were removed from analysis.

One female, “HUC”, unexpectedly left her social group for 9 days, then disappeared and is presumed dead. During her time alone, her activity decreased, and her HRV increased compared to before leaving her group (Fig. S2). See Case Study in Supporting Information for full results.

## Discussion

Measuring physiological metrics in free-ranging vertebrates is challenging, but physiologger systems with remote transmission capabilities can help (Gauld et al., 2023; Wild, Wikelski, et al., 2023). We combined a BLE-enabled physiologger with a LoRaWAN-enabled collar and a mobile LoRaWAN gateway to flexibly download fine-grained physiological data on heart rate, heart rate variability, activity, and body temperature from wild non-human primates over tens to hundreds of meters for up to 11 months. We did so without affecting the animals’ natural behaviors or re-capturing individuals and while consuming minimal power.

Our approach builds on prior work employing custom or human-derived cardiac monitors (Laske et al., 2021). Our ICMs were made by Medtronic, which to date have been deployed in 27 species, including wild American black bears (*Ursus americanus*), brown bears (*Ursus arctos*), grey wolves (*Canis lupus*), moose (*Alces alces*), maned wolves (*Chrysocyon brachyurus*), and southern elephant seals (*Mirounga leonina*; Laske et al., 2021). One limitation is that Medtronic requires a collaborative agreement to use its devices in animals (all other components of our system are available for purchase or open source). However, commercial cardiac monitors are becoming available for pets (e.g., Maven Health Sensors, https://maven.pet/) and livestock (Stygar et al., 2021). Modifying existing devices or creating custom devices that leverage BLE and LoRaWAN in tandem should be feasible for many researchers (Brandl et al., 2025; Wascher et al. 2011).

Another limitation of our system was range: the distance over which we could successfully download data was substantially shorter than other LoRaWAN systems in wildlife, which can transmit data over kilometers (Gauld et al., 2023; Parlin et al., 2025). Our range was constrained by the antenna and electronics used in the LoRaWAN collars; these were originally designed for a device with a smaller casing and could be better optimized for increased range. It should also be possible to increase range by elevating the LoRaWAN gateway on a permanent structure, like a tower (Ojo et al., 2021).

The data on HR and HRV we report here are, to our knowledge, the first-ever published for a free-ranging wild primate. The HR and body temperature values we recorded fall within the range of published data in captive baboons (Bert et al., 2013; Kuo et al., 2018; Brain & Mitchell, 1999; Nyakudya et al., 2012). However, we observed greater variability in body temperature, which may reflect differences in implant location (variation=8.2°C in our subcutaneous implant compared to 5.3°C in a peritoneal implant (Brain & Mitchell, 1999). Together, the system presented offers a valuable option to understand wildlife health, physiology, and responses to environmental change.

## Acknowledgements

We gratefully acknowledge the support of the National Institutes of Health for the data represented here, especially through R61/R33AG078470. Current support also comes from NIH R01AG071684, R01AG075914, and the Max Planck Institute for Evolutionary Anthropology (MPI-EVA). We thank Duke University, Princeton University, MPI-EVA, and the University of Notre Dame for financial and logistical support over the years. In Kenya, our research was approved by the Wildlife Research Training Institute, Kenya Wildlife Service, the National Commission for Science, Technology, and Innovation, and the National Environmental Management Authority. We also thank the University of Nairobi, the Kenya Institute of Primate Research, the National Museums of Kenya, the members of the Amboseli-Longido pastoralist communities, the Enduimet Wildlife Management Area, and Ker & Downey Safaris, for their cooperation. Particular thanks go to the Amboseli Baboon Project field team (R.S. Mututua, S. Sayialel, J.K. Warutere, I.L. Siodi, L. Musembei, and D.T. Kanai) and to T. Wango and V. Oudu for their assistance in Nairobi. The baboon project database is managed by J. Gordon and C. Broderick with recent assistance by W. Wilbur and N. Learn. Database design and programming are provided by K. Pinc. Insertable cardiac monitors were donated by Medtronic with programming support and training by N. Laske. We thank Jeanne Altmann for her foundational contributions to the Amboseli Baboon Project data set. For a complete set of acknowledgments, please visit http://amboselibaboons.nd.edu/acknowledgements/.

## Author contributions

E.S.P.: conceptualization, data curation, formal analysis, investigation, methodology, visualization, writing – original draft, writing – review and editing

C.R.A.: conceptualization, methodology, writing – review and editing

A.G.B.: investigation, writing – review and editing

E.K.: investigation, writing – review and editing

A.N.N.: investigation, writing – review and editing

P.S.: conceptualization, resources, software, writing – review and editing

A.T.: resources, software, writing – review and editing

P.L.: conceptualization, writing – review and editing

R.M.: conceptualization, methodology, writing – review and editing

P.A.I.: conceptualization, writing – review and editing

J.T.: data curation, funding acquisition, writing – review and editing

H.P.: conceptualization, funding acquisition, writing – review and editing

M.Y.A..: conceptualization, funding acquisition, methodology, investigation, writing – review and editing

S.C.A.: data curation, funding acquisition, writing – review and editing

T.v.D.: resources, methodology, software, writing – review and editing

T.G.L.: resources, methodology, writing – review and editing

E.A.A.: conceptualization, data curation, funding acquisition, investigation, project administration, writing – review and editing

## Statement on inclusion

Our study includes authors from the United States, Brazil, Germany, the Netherlands, and Kenya (the country where the fieldwork took place). This set of people includes all individuals who contributed to the conceptualization, data collection, and data analysis, and all authors reviewed and approved the manuscript.

## Data availability statement

Data and code required to recreate these analyses are available on github at https://github.com/e-s-person/lorawan_methods.

## References

Alberts, S. C., & Altmann, J. (2012). The Amboseli Baboon Research Project: 40 Years of Continuity and Change. In P. M. Kappeler & D. P. Watts (Eds.), Long-Term Field Studies of Primates (pp. 261–287). Springer. 10.1007/978-3-642-22514-7_12

Andreadis, C. R., Kulahci, I. G., Ndung’u, J., Kigen, D., Kimiti, P., Mugambi, K. R., Laske, N. R., Mwadime, J., Wanjala, N., Pontzer, H., Laske, T. G., Akinyi, M. Y., & Archie, E. A. (2026). Using insertable cardiac monitors to test determinants of heart rate and activity in captive baboons. bioRxiv. 10.64898/2026.03.13.710869

Antoine-Santoni, T., Gualtieri, J.-S., Manicacci, F.-M., & Aiello, A. (2018). AMBLoRa: A Wireless Tracking and Sensor System Using Long Range Communication to Monitor Animal Behavior. Proceedings of the Seventh International Conference on Smart Cities, Systems, Devices and Technologies, 35–40. https://hal.science/hal-03028480

Bert, A. A., Drake, W. B., Quinn, R. W., Brasky, K. M., O’Brien, J. E., Lofland, G. K., & Hopkins, R. A. (2013). Transesophageal echocardiography in healthy young adult male baboons (Papio hamadryas anubis): Normal cardiac anatomy and function in subhuman primates compared to humans. Progress in Pediatric Cardiology, 35(2), 109–120. 10.1016/j.ppedcard.2013.09.002

Brain, C., & Mitchell, D. (1999). Body Temperature Changes in Free-ranging Baboons (Papio hamadryas ursinus) in the Namib Desert, Namibia. International Journal of Primatology, 20(4), 585–598. Psychology Collection; Social Science Premium Collection. 10.1023/A:1020394824547

Brandl, H. B., Klarevas-Irby, J. A., Zuñiga, D., Hansen Wheat, C., Christensen, C., Omengo, F., Nzomo, C., Cherono, W., Nyaguthii, B., & Farine, D. R. (2025). The physiological cost of leadership in collective movements. Current Biology, 35(16), 4003–4010.e4. 10.1016/j.cub.2025.06.065

Brocaar, O. (2025). ChirpStack, open-source LoRaWAN® Network Server. v4. https://www.chirpstack.io/

Devalal, S., & Karthikeyan, A. (2018). LoRa Technology—An Overview. 2018 Second International Conference on Electronics, Communication and Aerospace Technology (ICECA), 284–290. 10.1109/ICECA.2018.8474715

Forin-Wiart, M.-A., Enstipp, M. R., Le Maho, Y., & Handrich, Y. (2019). Why implantation of bio-loggers may improve our understanding of how animals cope within their natural environment. Integrative Zoology, 14(1), 48–64. 10.1111/1749-4877.12364

Gaidica, M., & Dantzer, B. (2020). Quantifying the Autonomic Response to Stressors—One Way to Expand the Definition of “Stress” in Animals. Integrative and Comparative Biology, 60(1), 113–125. 10.1093/icb/icaa009

Gauld, J., Atkinson, P. W., Silva, J. P., Senn, A., & Franco, A. M. A. (2023). Characterisation of a new lightweight LoRaWAN GPS bio-logger and deployment on griffon vultures Gyps fulvus. Animal Biotelemetry, 11(1), 17. 10.1186/s40317-023-00329-y

Hawkes, L. A., Fahlman, A., & Sato, K. (2021). What is physiologging? Introduction to the theme issue, part 2. Philosophical Transactions of the Royal Society B: Biological Sciences, 376(1831), 20210028. 10.1098/rstb.2021.0028

Kuo, A. H., Li, C., Huber, H. F., Nathanielsz, P. W., & Clarke, G. D. (2018). Ageing changes in biventricular cardiac function in male and female baboons (Papio spp.). The Journal of Physiology, 596(21), 5083–5098. 10.1113/JP276338

Kutinsky, I., Duncan, A., Danforth, M. D., Murray, S., Napier, J., McCain, S., & Murphy, H. W. (2023). Surgical placement of implantable cardiac loop recorders in great apes. American Journal of Primatology, 85(3), e23471. 10.1002/ajp.23471

Laske, T. G., Garshelis, D. L., & Iaizzo, P. A. (2011). Monitoring the wild black bear’s reaction to human and environmental stressors. BMC Physiology, 11(1), 13. 10.1186/1472-6793-11-13

Laske, T. G., Evans, A. L., Arnemo, J. M., Iles, T. L., Ditmer, M. A., Fröbert, O., Garshelis, D. L., & Iaizzo, P. A. (2018). Development and utilization of implantable cardiac monitors in free-ranging American black and Eurasian brown bears: System evolution and lessons learned. Animal Biotelemetry, 6(1), 13. 10.1186/s40317-018-0157-z

Laske, T. G., Iaizzo, P. A., & Garshelis, D. L. (2017). Six Years in the Life of a Mother Bear— The Longest Continuous Heart Rate Recordings from a Free-Ranging Mammal. Scientific Reports, 7(1), 40732. 10.1038/srep40732

Laske, T. G., Garshelis, D. L., Iles, T. L., Iaizzo, P. A. (2021). An engineering perspective on the development and evolution of implantable cardiac monitors in free-living animals. Philosophical Transactions of the Royal Society B: Biological Sciences, 376(1830):20200217. 10.1098/rstb.2020.0217

Leimgruber, P., Songsasen, N., Stabach, J. A., Horning, M., Reed, D., Buk, T., Harwood, A., Layman, L., Mathews, C., Vance, M., Marinari, P., Helmick, K. E., Delaski, K. M., Ware, L. H., Jones, J. C., Silva, J. L. P., Laske, T. G., & Moraes, R. N. (2023). Providing baseline data for conservation–Heart rate monitoring in captive scimitar-horned oryx. Frontiers in Physiology, 14. 10.3389/fphys.2023.1079008

LoRa Alliance™ Strategy Committee. (2018). LoRaWANTM Geolocation Whitepaper. [White paper]. LoRa AllianceTM. https://resources.lora-alliance.org/whitepapers/lora-alliance-geolocation-whitepaper

Moraes, R. N., Laske, T. G., Leimgruber, P., Stabach, J. A., Marinari, P. E., Horning, M. M., Laske, N. R., Rodriguez, J. V., Eye, G. N., Kordell, J. E., Gonzalez, M., Eyring, T., Lemons, C., Helmick, K. E., Delaski, K. M., Ware, L. H., Jones, J. C., & Songsasen, N. (2021). Inside out: Heart rate monitoring to advance the welfare and conservation of maned wolves (Chrysocyon brachyurus). Conservation Physiology, 9(1), coab044. 10.1093/conphys/coab044

Nyakudya, T. T., Fuller, A., Meyer, L. C. R., Maloney, S. K., & Mitchell, D. (2012). Body Temperature and Physical Activity Correlates of the Menstrual Cycle in Chacma Baboons (Papio hamadryas ursinus). American Journal of Primatology, 74(12), 1143–1153. 10.1002/ajp.22073

Ojo, M. O., Adami, D., & Giordano, S. (2021). Experimental Evaluation of a LoRa Wildlife Monitoring Network in a Forest Vegetation Area. Future Internet, 13(5), 115. 10.3390/fi13050115

Parlin, A. F., Horning, N. A., Alstad, J. P., Cosentino, B. J., & Gibbs, J. P. (2025). Low-cost, LoRa GNSS tracker for wildlife monitoring. HardwareX, 23, e00669. 10.1016/j.ohx.2025.e00669

Perea, A. R., Rahman, S., Chen, H., Cox, A., Nyamuryekung’e, S., Bakir, M., Cao, H., Estell, R., Bestelmeyer, B., Cibils, A. F., & Utsumi, S. (2025). Use of Lora Wan Wireless Sensor Data Transmission and Machine Learning Models to Classify the Behavior of Beef Cows Grazing Desert Rangelands in the Southwest United States (SSRN Scholarly Paper No. 5149569). Social Science Research Network. 10.2139/ssrn.5149569

Stygar, A.H., Gómez, Y., Berteselli, G.V., Dalla Costa, E., Canali, E., Niemi, J.K., Llonch, P., & Pastell, M., (2021). A Systematic Review on Commercially Available and Validated Sensor Technologies for Welfare Assessment of Dairy Cattle. Frontiers in Veterinary Science, 8. 10.3389/fvets.2021.634338

Vilgalys, T. P., Fogel, A. S., Anderson, J. A., Mututua, R. S., Warutere, J. K., Siodi, I. L., Kim, S. Y., Voyles, T. N., Robinson, J. A., Wall, J. D., Archie, E. A., Alberts, S. C., & Tung, J. (2022). Selection against admixture and gene regulatory divergence in a long-term primate field study. Science, 377(6606), 635–641. 10.1126/science.abm4917

Wascher, C. A. F., Weiß, B. M., Arnold, W., & Kotrschal, K. (2011). Physiological implications of pair-bond status in greylag geese. Biology Letters, 8(3), 347–350. 10.1098/rsbl.2011.0917

Wascher, C. A. F. (2021). Heart rate as a measure of emotional arousal in evolutionary biology. Philosophical Transactions of the Royal Society B: Biological Sciences, 376(1831), 20200479. 10.1098/rstb.2020.0479

Wild, T. A., van Schalkwyk, L., Viljoen, P., Heine, G., Richter, N., Vorneweg, B., Koblitz, J. C., Dechmann, D. K. N., Rogers, W., Partecke, J., Linek, N., Volkmer, T., Gregersen, T., Havmøller, R. W., Morelle, K., Daim, A., Wiesner, M., Wolter, K., Fiedler, W., … Wikelski, M. (2023). A multi-species evaluation of digital wildlife monitoring using the Sigfox IoT network. Animal Biotelemetry, 11(1), 13. 10.1186/s40317-023-00326-1

Wild, T. A., Wikelski, M., Tyndel, S., Alarcón-Nieto, G., Klump, B. C., Aplin, L. M., Meboldt, M., & Williams, H. J. (2023). Internet on animals: Wi-Fi-enabled devices provide a solution for big data transmission in biologging. Methods in Ecology and Evolution, 14(1), 87–102. 10.1111/2041-210X.13798

Williams, H. J., Shipley, J. R., Rutz, C., Wikelski, M., Wilkes, M., & Hawkes, L. A. (2021). Future trends in measuring physiology in free-living animals. Philosophical Transactions of the Royal Society B: Biological Sciences, 376(1831), 20200230. 10.1098/rstb.2020.0230

