## Supporting Information for "Real-time heart rate in the wild: remote collection of cardiac data in baboons using a low-power Bluetooth and LoRaWAN system"

### Supplementary methods

ICM insertion. Subjects were darted and anesthetized with a handheld blowgun loaded with a Tiletamine-Zolazepam-bearing dart, using a minimally invasive protocol used by the ABRP (Altmann et al., 1996; Tung et al., 2015). To minimize disruption to the group, darting only occurred when no baboons would observe the dart delivery, and no more than two animals were darted per day. Anesthetized baboons were quickly removed to a processing site distant from the rest of the group and monitored for the duration of the procedure.

To insert the ICM, the skin over the left pectoral muscle was shaved and cleaned with ethanol and betadine povidone-iodine solution. We made a 0.5 cm incision perpendicular to the insertion area and used the tools provided by Medtronic to create a ~7 cm subcutaneous pocket into which we inserted the ICM. The incision was closed with two stitches and we applied betadine povidone-iodine ointment on the incision site. Each animal was given long-lasting antibiotic via intramuscular injection (Penicillin-Streptomycin, 1cc/10kg body mass).

Following implantation, each animal was fitted with a Smart Parks collar paired to the individual's ICM. Baboons recovered in a draped holding cage (3-4 h) and were then released in an area where they had full view of their social group. All subjects rejoined their groups quickly and without incident. The animals were monitored every 1-3 days by ABRP observers who looked for signs of infection or illness; none were observed in any subject.

<u>Summary of physiological data.</u> For each subject, we calculated mean HR, HRV, activity score, and body temperature for each day during daytime hours when baboons are most active (06:30-18:30, approximately sunrise to sunset) and nighttime hours when baboons are in "sleeping trees" and less active (18:30-06:30). We also calculated a mean "resting" HR, HRV, activity, and body temperature as the mean of the three 5-minute values observed between 05:00-05:15 when HR and activity were typically at their lowest point (Speed et al., 2023). Resting HR and HRV are often linked to health outcomes in humans and animal models (Carnevali & Sgoifo, 2014; Johansen et al., 2013) and may reflect chronic stress (Jiryis et al., 2022).

<u>Case study.</u> We analyzed individual HR, HRV, activity, and body temperature from a female baboon (HUC) who left her social group for unknown reasons and stayed alone for 9 days in August 2025, six months after implantation of the ICM. HUC was four months pregnant at the time of her disappearance; pregnancy was known from well-validated signs of pregnancy in baboons and near-daily monitoring of reproductive state by ABRP observers (S. A. Altmann, 1973; Gesquiere et al., 2018). At the time of implantation and prior to her disappearance in August 2025, HUC appeared to be in good health, although at 22 years of age she was relatively old for a female baboon in Amboseli. Only 3.5% of female baboons (22 of 592 reported in Bronikowski et al., 2016) born in the population reach age 22 years.

We estimated the impact of living alone on each of HUC's physiological metrics using a Bayesian causal impact analysis implemented in the 'CausalImpact' R package (v 1.4.1;

Broderson et al., 2015). For HR, HRV, activity, and body temperature, we used 19 days of data prior to HUC's departure (1-19 Aug 2025) to inform a time series model. We then created a counterfactual prediction of her expected physiological metrics had she remained with her group for the 9 days she was alone (21-30 Aug 2025; data from 20 August are excluded because HUC's group was not observed that day). We compared the observed metrics to the counterfactual prediction to infer the difference in each metric linked to HUC's period of social isolation. For each model, we included mean hourly air temperature recorded from a weather station (ClimaVue 50 Digital Weather Sensor, Campbell Scientific, Logan, UT, USA) as a covariate. For the model of HRV, we also included HR as a covariate because these values are related (Sacha, 2014).

We also visualized changes in HUC's HR, HRV, activity, and body temperature during the 9 days she was separated from her group compared to the 19 days prior to her separation, including: (1) 5-minute median HR during the day, night, and while resting (05:00-05:15); (2) 5-minute HRV during the day, night, and while resting; (3) 5-minute activity scores during the day; (4) sleep disruption, calculated as the mean number of "high activity" minutes per night (high activity=5-minute activity scores >100 on Medtronic's unitless scale from 0 to 255); (5) 1-hour body temperatures during the day and at night. Analyses and figures were performed in R (v. 4.4.3; R Core Team, 2025)

**Supplementary results**

<u>Case study.</u> On August 21$^{st}$ 2025, HUC was observed missing from her group during monitoring. After a search, she was located several kilometers away, foraging and resting alone. She appeared uninjured and was not visibly ill, but such behavior is consistent with the profile of an animal who is in poor health and close to death (HUC was 22.7 years old at the time of her disappearance). After 9 days of separation from her group, we were unable to detect her collar

and she could not be located during a search of known sleeping sites, so she was presumed dead. Our failure to detect signals from HUC's collar could have been caused by her moving outside our range of detection, or by damage to her collar during a predator attack or other incident. Several months after her disappearance, her collar was recovered ~1 km from the site she was last seen. The collar was heavily damaged, likely by a human or predator though it is unknown if this damage was caused around the time of her disappearance and/or death.

During the 9-day period of separation, HUC's HR increased slightly with an observed mean of 87.3 bpm compared to the expected mean of 85.3 bpm in the counterfactual prediction, although this effect was not statistically significant (causal impact analysis: $P$=0.462; Fig. S2A). HUC's HR also appeared more variable when she was alone, and her median resting HR was higher, which can be an indication of stress (visualization: Fig. S2B; Johansen et al., 2013). Contrary to our expectations, HUC showed significantly higher HRV during this period of isolation, with an observed mean HRV of 7.2 ms compared to the expected mean of 6.5 ms (causal impact analysis: $P$=0.048; Fig. S2C,D). Although activation of the stress response is expected to reduce HRV (Kim et al., 2018), HUC's mean activity decreased 17% compared to her predicted activity (causal impact analysis: $P$=0.041; Fig. S2E,F). We also found some evidence for disrupted sleep: HUC showed on average 15 more minutes of "high activity" (scores >100) at night when she was alone compared to when she was with her group (visualization: Fig. S2G). We did not observe any significant change in her body temperature relative to the predicted mean.

### Discussion

HUC's separation from her social group provided a useful case study to highlight the value of real-time physiological data. It is highly unusual for female baboons in Amboseli to leave their social groups unless they are physically unable to keep up. Although HUC appeared uninjured, she never rejoined her group. Thus, it is likely she had an injury or illness we could not see that left her unable to stay with her group. Her daytime activity was substantially lower during her period of isolation than when she was with the group, so she may have been restricting her movement or resting more than usual. She was also four months pregnant during this time, which could have exacerbated any underlying issues (J. Altmann, 1983; Ellison, 2003; Gesquiere et al., 2024). Additionally, she showed higher activity at night, which could indicate a need for increased watchfulness for predators or disturbed sleep due to stress or lack of opportunities for thermoregulation via huddling (Pawlyk et al., 2008).

Regardless of the reason for the separation, isolation is often stressful for social animals (Mumtaz et al., 2018). Hence, we predicted an increase in HR and a decrease in HRV during this period as both reflect stress in humans and other animals (Grippo et al., 2007; Gaidica & Dantzer, 2020; Kim et al., 2018). However, we did not observe a significant increase in HR and we were surprised to find that HUC's HRV was higher during the isolated period, at times reaching an order of magnitude greater than her typical mean. We can think of at least three explanations for these patterns. First, HUC's high HRV may be pathological. In humans, abnormally high HRV can indicate underlying problems like heart arrhythmias and more generally can be associated with poor health outcomes (Bruyne et al., 1999; Stein et al., 2005). Second, HRV has been shown to vary throughout pregnancy in humans, with increases observed in the third trimester and during labor (Musa et al., 2017; Sarhaddi et al., 2022). It is possible that physiological changes due to pregnancy or a fetal loss contributed to the unusual HRV values we observed (fetal losses are well-documented and common in this population, accounting for 14% of pregnancies after pregnancy was detectable by observers; Fogel et al. 2022). Third, HUC may have experienced considerable psychosocial stress while trying to remain with her group, and this stress was alleviated once she left her group. In support of this possibility, research on vulturine guineafowl shows that HRV is reduced when individual movement decisions conflict with those of their social group (Brandl et al., 2025).

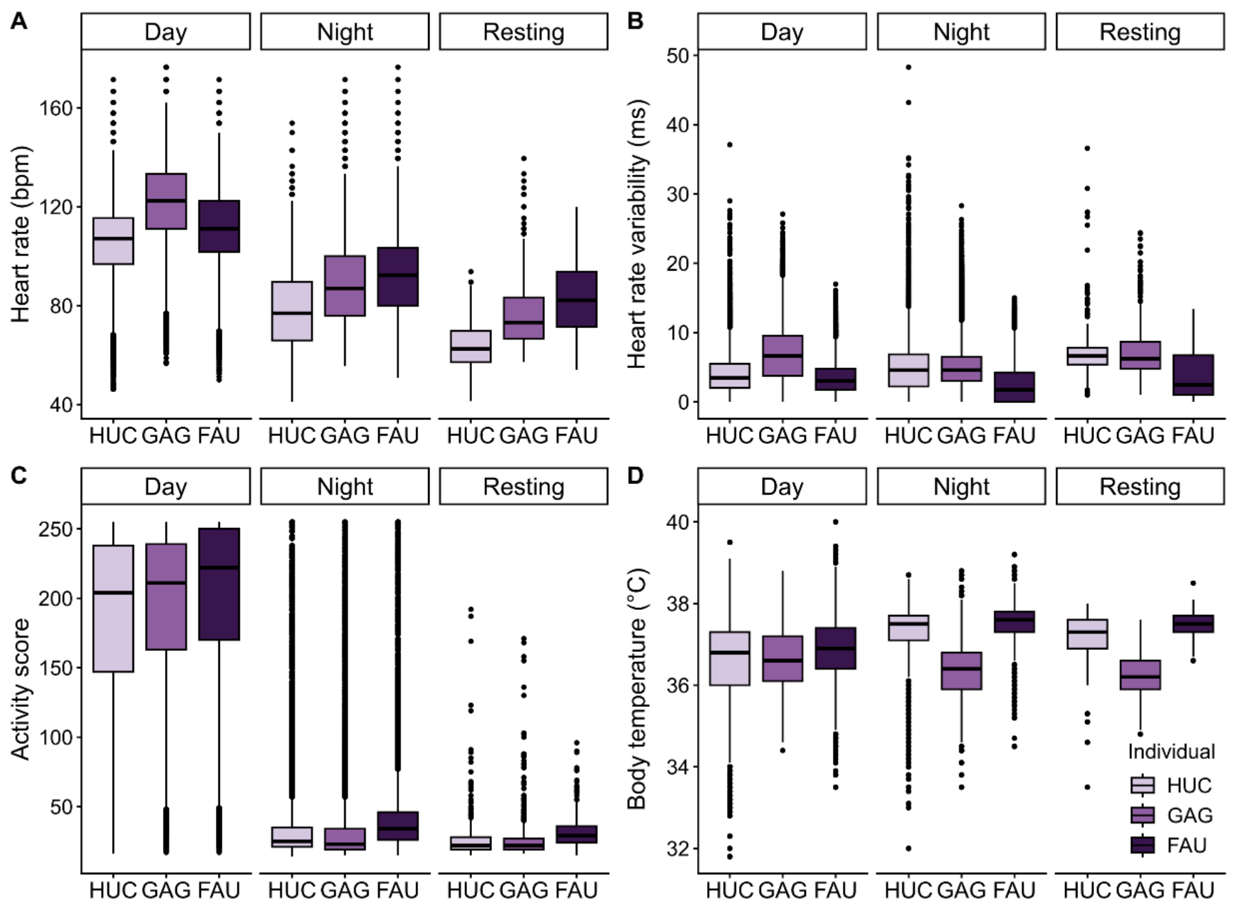

**Figure S1. Median and interquartile ranges for (A) Heart rate, (B) heart rate variability, (C) activity score, (D) body temperature from three female baboons over six months (March-August 2025).** Data are divided into three time periods: day (06:30-18:30), night (18:30-06:30) and the designated resting period (05:00-05:15). In each box plot, the horizontal bars represent medians; box tops and bottoms represent the 25th and 75th percentiles; whiskers are 1.5 times the interquartile range; dots are data outside the interquartile range.

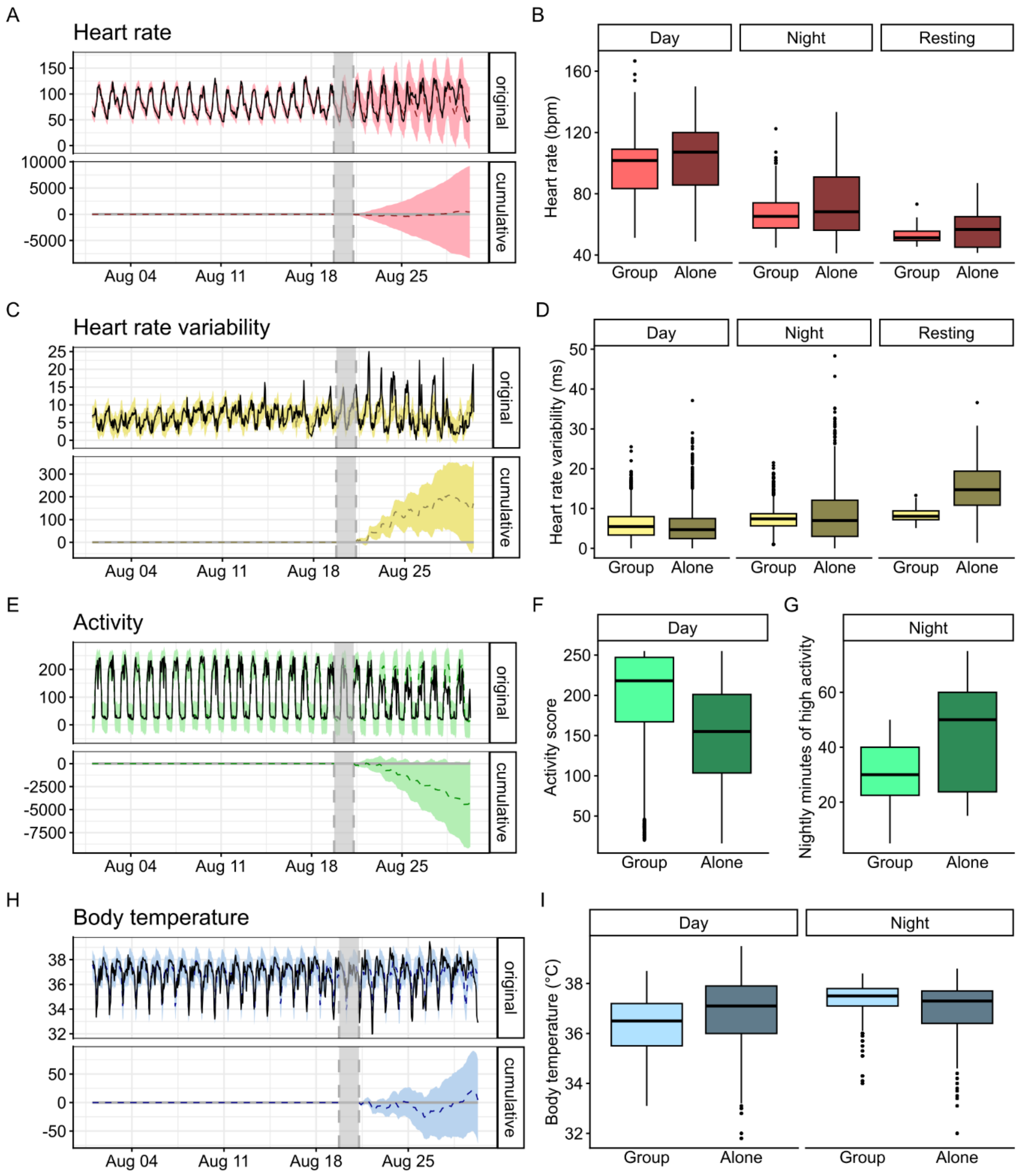

**Figure S2. Baboon physiology during a period of social isolation**. Plots in the left column (A, C, E, H) are time series illustrating change in each physiological metric attributed to HUC's social isolation. The solid black line in the top plot of each time series shows the observed data, while the dashed line shows the counterfactual prediction of the expected data had HUC not left her social group. Note that the dashed line is not visually apparent throughout much of the 9 day period of isolation because it falls close to the solid black (observed) line. The colored areas show the 95% confidence interval of the predictions. The bottom plot of each time series shows the cumulative difference between the observed and predicted data. The grey shaded bar in each plot indicates the period where HUC's group status was unknown (these data were excluded from the model); data to the left of the shaded area show HUC with her social group; data to the right of the shaded area HUC was separated from her group. Box plots in the right column (B, D, F, I) indicate median and interquartile ranges for each measure during the period HUC was with her group and when she was alone. (A and B) HR, (C and D) HRV, (E) activity score, (F) daytime activity only, (G) nighttime minutes of high activity, (H and I) body temperature.

### References

Altmann, J. (1983). Costs of reproduction in baboons. In *Behavioral Energetics: The cost of survival in vertebrates* (pp. 67–88). W.P. Aspey and S.I. Lustick. Columbus, Ohio State University Press.

Altmann, J., Alberts, S. C., Haines, S. A., Dubach, J., Muruthi, P., Coote, T., Geffen, E., Cheesman, D. J., Mututua, R. S., Saiyalel, S. N., Wayne, R. K., Lacy, R. C., & Bruford, M. W. (1996). Behavior predicts genes structure in a wild primate group. Proceedings of the National Academy of Sciences, 93(12), 5797–5801. https://doi.org/10.1073/pnas.93.12.5797

Altmann, S. A. (1973). The Pregnancy Sign in Savannah Baboons. *The Journal of Zoo Animal Medicine*, *4*(2), 8–12. https://doi.org/10.2307/20094180

Bronikowski, A. M., Cords, M., Alberts, S. C., Altmann, J., Brockman, D. K., Fedigan, L. M., Pusey, A., Stoinski, T., Strier, K. B., & Morris, W. F. (2016). Female and male life tables for seven wild primate species. *Scientific Data*, *3*(1), 160006. https://doi.org/10.1038/sdata.2016.6

Broderson, K. H., Gallusser, F., Koehler, J., Remy, N., & Scott, S. L. (2015). Inferring causal impact using Bayesian structural time-series models. *The Annals of Applied Statistics, 9*(1): 247-274. https://doi.org/10.1214/14-AOAS788

Bruyne, M. C. de, Kors, J. A., Hoes, A. W., Klootwijk, P., Dekker, J. M., Hofman, A., van Bemmel, J. H., & Grobbee, D. E. (1999). Both Decreased and Increased Heart Rate Variability on the Standard 10-Second Electrocardiogram Predict Cardiac Mortality in the Elderly: The Rotterdam Study. *American Journal of Epidemiology*, *150*(12), 1282–1288. https://doi.org/10.1093/oxfordjournals.aje.a009959

Carnevali, L., & Sgoifo, A. (2014). Vagal modulation of resting heart rate in rats: The role of stress, psychosocial factors, and physical exercise. Frontiers in Physiology, 5. https://doi.org/10.3389/fphys.2014.00118

Ellison, P. T. (2003). Energetics and reproductive effort. *American Journal of Human Biology*, *15*(3), 342–351. https://doi.org/10.1002/ajhb.10152

Fogel, A. S., Oduor, P. O., Nyongesa, A. W., Kimwele, C. N., Alberts, S. C., Archie, E. A., & Tung, J. (2023). Ecology and age, but not genetic ancestry, predict fetal loss in a wild baboon hybrid zone. *American Journal of Biological Anthropology, 180*(4), 618-632. https://doi.org/10.1002/ajpa.24686

Gesquiere, L. R., Altmann, J., Archie, E. A., & Alberts, S. C. (2018). Interbirth intervals in wild baboons: Environmental predictors and hormonal correlates. *American Journal of Physical Anthropology*, *166*(1), 107–126. https://doi.org/10.1002/ajpa.23407

Gesquiere, L. R., Adjangba, C., Wango, T. L., Oudu, V. K., Mututua, R. S., Warutere J. K., Siodi, I. L., Campos, F. A., Archie, E. A., Markham, A. C., & Alberts, S. C. (2024). Thyroid hormone concentrations in female baboons: Metabolic consequences of living in a highly seasonal environment. *Hormones and Behavior*. *161*, 105505. https://doi.org/10.1016/j.yhbeh.2024.105505

Jiryis, T., Magal, N., Fructher, E., Hertz, U., & Admon, R. (2022). Resting-state heart rate variability (HRV) mediates the association between perceived chronic stress and ambiguity avoidance. Scientific Reports, 12(1), 17645. https://doi.org/10.1038/s41598-022-22584-4

Johansen, C. D., Olsen, R. H., Pedersen, L. R., Kumarathurai, P., Mouridsen, M. R., Binici, Z., Intzilakis, T., Køber, L., & Sajadieh, A. (2013). Resting, night-time, and 24 h heart rate as markers of cardiovascular risk in middle-aged and elderly men and women with no apparent heart disease. European Heart Journal, 34(23), 1732–1739. https://doi.org/10.1093/eurheartj/ehs449

Kim, H.-G., Cheon, E.-J., Bai, D.-S., Lee, Y. H., & Koo, B.-H. (2018). Stress and Heart Rate Variability: A Meta-Analysis and Review of the Literature. *Psychiatry Investigation*, *15*(3), 235–245. https://doi.org/10.30773/pi.2017.08.17

Mumtaz, F., Khan, M. I., Zubair, M., & Dehpour, A. R. (2018). Neurobiology and consequences of social isolation stress in animal model—A comprehensive review. *Biomedicine & Pharmacotherapy*, *105*, 1205–1222. https://doi.org/10.1016/j.biopha.2018.05.086

Musa, S. M., Adam, I., Hassan, N. G., Rayis, D. A., & Lutfi, M. F. (2017). Maternal Heart Rate Variability during the First Stage of Labor. *Frontiers in Physiology*, *8*(774). https://doi.org/10.3389/fphys.2017.00774

Pawlyk, A. C., Morrison, A. R., Ross, R. J., & Brennan, F. X. (2008). Stress-induced changes in sleep in rodents: Models and mechanisms. *Neuroscience & Biobehavioral Reviews*, *32*(1), 99–117. https://doi.org/10.1016/j.neubiorev.2007.06.001

R Core Team. (2025). R: A language and environment for statistical computing. [Computer software]. R Foundation for Statistical Computing, Vienna, Austria. https://www.R-project.org

Sacha, J. (2014). Interaction between Heart Rate and Heart Rate Variability. *Annals of Noninvasive Electrocardiology*, *19*(3), 207–216. https://doi.org/10.1111/anec.12148

Sarhaddi, F., Azimi, I., Axelin, A., Niela-Vilen, H., Liljeberg, P., & Rahmani, A. M. (2022). Trends in Heart Rate and Heart Rate Variability During Pregnancy and the 3-Month Postpartum

Period: Continuous Monitoring in a Free-living Context. *JMIR mHealth and uHealth*, *10*(6), e33458. https://doi.org/10.2196/33458

Speed, C., Arneil, T., Harle, R., Wilson, A., Karthikesalingam, A., McConnell, M., & Phillips, J. (2023). Measure by measure: Resting heart rate across the 24-hour cycle. PLOS Digital Health, 2(4), e0000236. https://doi.org/10.1371/journal.pdig.0000236

Stein, P. K., Domitrovich, P. P., Hui, N., Rautaharju, P., & Gottdiener, J. (2005). Sometimes Higher Heart Rate Variability Is Not Better Heart Rate Variability: Results of Graphical and Nonlinear Analyses. *Journal of Cardiovascular Electrophysiology*, *16*(9), 954–959. https://doi.org/10.1111/j.1540-8167.2005.40788.x

Tung, J., Zhou, X., Alberts, S. C., Stephens, M., & Gilad, Y. (2015). The genetic architecture of gene expression levels in wild baboons. eLife, 4, e04729. https://doi.org/10.7554/eLife.04729
